# Exploratory bioinformatics analysis reveals importance of “junk” DNA in early embryo development

**DOI:** 10.1101/079921

**Authors:** Steven Xijin Ge

## Abstract

**Background:** Instead of testing predefined hypotheses, the goal of exploratory data analysis (EDA) is to find what data can tell us. Following this strategy, we re-analyzed a large body of genomic data to investigate how the early mouse embryos develop from fertilized eggs through a complex, poorly understood process.

**Results:** Starting with a single-cell RNA-seq dataset of 259 mouse embryonic cells from zygote to blastocyst stages, we reconstructed the temporal and spatial dynamics of gene expression. Our analyses revealed similarities in the expression patterns of regular genes and those of retrotransposons, and the enrichment of transposable elements in the promoters of corresponding genes. Long Terminal Repeats (LTRs) are associated with transient, strong induction of many nearby genes at the 2-4 cell stages, probably by providing binding sites for Obox and other homeobox factors. The presence of B1 and B2 SINEs (Short Interspersed Nuclear Elements) in promoters is highly correlated with broad upregulation of intracellular genes in a dosage-and distance-dependent manner. Such enhancer-like effects are also found for human Alu and bovine tRNA SINEs. Promoters for genes specifically expressed in embryonic stem cells (ESCs) are rich in B1 and B2 SINEs, but low in CpG islands.

**Conclusions:** Our results provide evidence that transposable elements may play a significant role in establishing the expression landscape in early embryos and stem cells. This study also demonstrates that open-ended, exploratory analysis aimed at a broad understanding of a complex process can pinpoint specific mechanisms for further study.

**Major finding:**

- Single-cell RNA-seq data enables estimation of retrotransposon expression during PD
- Similar expression dynamics of retrotransposons and regular genes during PD
- Long terminal repeats may be essential for the 1^st^ wave of gene expression
- Obox homeobox factors are possible regulators of PD, upstream of Zscan4
- SINE repeats predict expression of nearby genes in murine, human and bovine embryos
- Exploratory analysis of large single-cell data pinpoints developmental pathways

## Introduction

In the initial stage of mammalian development, zygotes undergo maternal-to-zygotic transition (MZT)^1^, during which maternal factors are eliminated while mRNA and protein synthesis using zygotic genome are initiated to take control of embryo development. A highly coordinated cascade of regulatory mechanisms unfolds rapidly to give rise to several cell lineages and the formation of blastocysts^2^. Understanding the complex process of pre-implantation development (PD) is important for both fertility related interventions as well as manipulation of embryonic stem cells (ESCs).

Many gene expression studies of the early mouse embryos has been carried out using expressed sequence tags (ESTs)^3–6^, DNA microarrays^7–11^, RNA-sequencing (RNA-seq)^12^, and, more recently, single-cell RNA-seq^13–15^. These studies documented the dynamic waves of gene expression via the regulation of thousands of genes at different stages. Using the powerful single-cell RNA-seq technique^14,16^, Deng *et al.* analyzed hundreds of cells from embryos of mixed background mice and found strong evidence for random and widespread monoallelic expression^14^. With sequence-level detail at single-cell resolution, this and other similar datasets^13,15,17^ can be used for in-depth study of gene regulation during PD. The epigenetic remodeling of maternal and paternal genomes was also revealed by DNA methylation profiling^18–20^. In Zebra fish, transcription factors (TFs) such as POU5F1, NANOG, and SOXB1 were found to activate zygotic gene expression^2,21^. But the molecular mechanisms of PD remain poorly understood.

Retrotransposon expression is a defining event in genome reprogramming during PD^22^. Transposable elements (TEs) cover 30-50% of mammalian genomes^23^. Some are actively transcribed^24,25^, even retrotransposed, as much of the genome is briefly hypomethylated in early embryos. The expression of retrotransposons is dynamic and stage-specific^20,24,26–28^. Evsikov et al. showed that retrotransposons, especially long terminal repeats (LTRs), are abundantly represented in mouse embryo at 2-cell stage ^25^. Class III LTRs such as MERV-L family LTRs are transcribed at extremely high levels, accounting for about 3% of total transcriptional output at this stage^29^. Human endogenous retroviruses (ERVs) HERVK LTR are expressed at zygotic genome activation (ZGA) at eight-cell stage^26^. Long interspersed elements (LINEs), another major type of retrotransposon, are also expressed during PD^30^, similar to short interspersed nuclear elements (SINEs). Interestingly, some endogenous retroviral activities have been found to be associated with and can serve as markers of pluripotency and totipotency^27,28,31^ in stem cell populations, underlying the importance of study retrotransposon expression.

Transcription of retrotransposons can directly influence the expression of neighboring genes. By analyzing EST libraries, Peaston *et al.* analyzed the expression of different types of TEs during PD and found that TEs, especially MERV-L family LTRs, provide alternative 5’ first exons to 41 chimeric transcripts in 2-cell mouse embryos^32^. Besides LTRs, long interspersed elements (LINEs) and short interspersed nuclear elements (SINEs) also contributed a small number of chimeric transcripts. LINEs have also been found to initiate fusion transcripts in other studies^33^. More broadly, by analyzing cap-selected 5’ end of mouse and human transcripts from various embryonic and adult tissues, Faulkner et al.^24^ found that 6-30% of all transcripts initiates from TEs^24^. These transcripts are often tissue-specific. Studies on long noncoding RNAs (lncRNAs) also identified ~30,000 TEs critical for the biogenesis of about 30% of total lncRNAs sequences^34^.

TEs can also influence gene expression indirectly by shaping the epigenetics landscape^18–20,35–37^, as GC-rich SINEs tend to be found near genes while LINEs has the opposite distribution. Some transcripts from TEs give rise to small RNAs that involve in post-transcriptional regulation during PD^38–40^. LINE-1 RNA can regulate its own expression^20^. Chip-Seq data in human cells shows that TEs contribute 25% of binding sites for transcription factors (TFs) critical for embryonic stem cells such as POU5F1, NANOG and CTCF^34^. Xie et al. ^9^ compared expression dynamics of PD in human, mouse and bovine and found substantial difference in co-expression network, which could be partly due to the cis-regulatory modules provided by species-specific transposons. Recently, Tӧhӧnen *et al.* used single-cell RNA-seq to analyze 348 single-cells from early human development^15^. They found that the promoters of 32 genes upregulated early at 4-cell stage are enriched with Alu elements that harbor motifs, including a “TAATCC” core motif bound by PITX and OTX homeobox family^15^. These studies provide substantial evidence for TEs’ broad role in regulating gene expression during PD.

In this study, we use exploratory data analysis (EDA) approach to study the dynamic expression of both regular genes and TEs in mouse PD. First proposed by Tukey, the goal of EDA^41^ is to ***explore the data and find what it can tell us.*** It is a philosophically different approach to statistical analysis, not a new set of tools. In addition to data visualization, all statistical techniques can be used to assess distributions, structures and dependencies that are useful for modeling^42,43^. More importantly, EDA can reveal important characteristics and trends, which can help formulate new hypotheses for further investigation^41^.

The open-ended EDA approach can serve as a general strategy to generate new hypotheses from biomedical data repositories. Such inductive methods can complement hypothesis-driven approaches to form an iterative process of ongoing research^44,45^. Our knowledge about many fundamental biological processes remains limited and fragmented. We can take advantage of the massive genome-wide datasets to learn about biological systems, akin to the study of an unknown planet in space exploration. The main challenge is to organically combine multiple datasets and analytical tools to gain actionable insights into complex biological processes.

This study represents an attempt of exploratory bioinformatics analysis (EBA) on gene regulation in PD. Starting from the large single-cell RNA-seq data of Deng et al.^14^, *our approach is to systematically observe the gene expression dynamics to help develop a broad understanding of the regulatory mechanisms of this complex process*. Our goal is to produce insights that can lead to novel, testable hypotheses on early embryo development. The data was analyzed alongside other expression and epigenetic studies of PD, using a various tools and annotation databases (See flowchart in Figure 1A). Our analyses provide evidence for co-regulation of transposons and regular genes in early embryo development in mouse, followed up with similar observation in human, bovine and zebrafish. This adds to existing evidence that the expansion of species-specific TEs may help rewire developmental pathways^34,46^. Motif analysis of promoter sequences of stage-specific genes identified many homeobox domain TFs as potential regulators of PD, which could be experimentally tested. We also examined the non-random distribution of TEs in the mouse genome and the enrichment of SINEs in the promoters of genes specifically expressed in embryonic stem cells (ESCs). Finally, we discussed the evolutionary benefits of transposons in promoting genetic diversity, especially in slow-reproducing animals.

**Figure 1.**
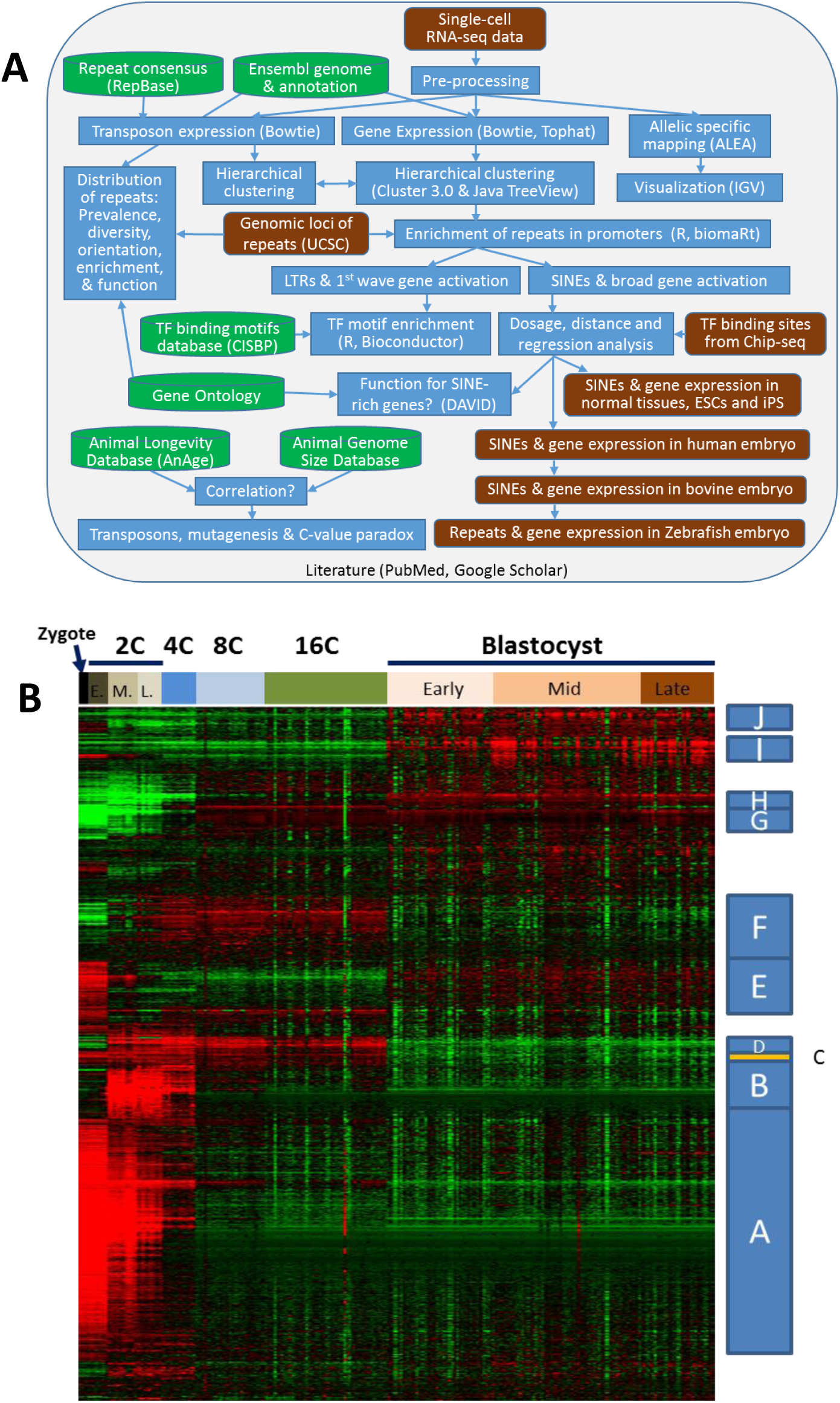
A) Exploratory bioinformatics investigation on gene regulation in mouse pre-implantation development. B) Hierarchical clustering of gene expression during PD. Each of the 12,000 rows represents a gene. Columns correspond to samples labeled by developmental stages (E: early, M: middle, and L: late). Red indicates expression levels higher than average for the row. Expression lower than average is shown in green.

## Results

To study allelic-specific transcription, Deng *et al.*^14^ analyzed individual cells from mixed lineage mouse embryos (CAST/EiJ females mated with C57BL/6 male) using the single-cell RNA-seq technique^47^. Excluding technical control samples, 259 RNA-seq libraries were generated, representing cells from zygote to blastocyst stages. The 2-cell (2C) stage is divided into early, middle (mid) and late phases. Each library contains about 22 million reads, mostly 43 base pair (bp) long. With a total of about 6 billion reads, this massive data enables in-depth EBI on the temporal and spatial regulation of transcription during PD.

### Similar expression patterns for regular genes and retrotransposons

The raw sequence reads were re-analyzed to quantify gene expression using Tophat and Cufflinks programs^48^ with updated genome annotation from Ensembl^49^. Hierarchical clustering was used to analyze the expression pattern of 12,000 genes with sufficient change in expression across 259 individual cells at various developmental stages. In addition to temporal dynamics, this data can also reveal variability among cells at the same stage. Ten gene clusters were defined according to the similarity in temporal expression patterns (Figure 1B), similar to previous efforts based on DNA microarray^7^. See supplementary document for the gene lists and details. Cluster A includes 3310 genes highly expressed in oocytes and quickly reduced at 2-cell (2C) and 4-cell (4C) stages. These are mostly maternal mRNAs undergoing degradation, which is evident from allele-specific mapping (supplementary Figures S1-S4). Zygotic genome activation (ZGA) occurs during the 2C stage, evidenced by marked changes in gene expression between mid and early 2C stages (supplementary Figure S5). The 777 transcripts in cluster B are exclusively expressed at 2C and 4C stages. Cluster D genes are induced at mid 2C, but their transient expression lasts until the 16-cell (16C) stage. Cluster F genes are gradually upregulated at the 2C stage and downregulated in the blastocyst. Genes in clusters G to J are activated at various stages and remain highly expressed. Our goal is to find the regulatory mechanism behind these different patterns of expression.

During PD, much of the genome is de-methylated and transposons are actively transcribed. Previous studies have shown that their expression patterns are dynamic and stage-specific^20,24,26,27^. To estimate their abundance, we used TETranscripts software^50^ with a special index file derived from RepeatMasker data available at UCSC genome browser website^51^. Supplementary Figure S6 shows the expression pattern of some of the highly transcribed transposons. Supplementary Table S7 contains the detailed expression levels of all TEs. The most highly expressed is a retrovirus-like element MT-int (RepBase ID: MTAI), a long terminal repeat (LTR) of the ERVL-MaLR family. It is expressed from the zygote to the 4C stage, similar to Cluster A genes. MERV-L (RepBase ID: MT2_Mm) is an ERVL family LTR that is sharply induced by more than 500-fold at mid 2C before decreasing to low levels at 8C, showing an expression pattern similar to cluster B genes. This is in agreement with a previous report that MERV-L accounts for about 3% of total transcriptional output at this stage^29^. There are several other LTR elements with this type of expression, including MT2C_Mm, ORR1A2, ORR1A3, and MT2B2. Intracisternal A particle (IAP) elements are also transcribed between 2C and 16C as expected^52^, similar to cluster D genes. Besides LTRs, LINEs and SINEs are also expressed as expected. SINE transcripts increase modestly at the 2C stage and remain at that level through the blastocyst stage, similar to Cluster F and G genes. Retrotransposons are transcribed in a highly regulated manner during PD, with expression patterns mirroring those of regular genes.

Some MERV-L elements seem to give rise to a microRNA (miR-1194) from several genomic loci (Gm23215, Gm23551, Gm23943, Gm24617, Gm25042, and Gm26475). Low level expression of miR-1194 in ESCs has been detected^53^. Further study is needed to determine whether this microRNA aids the clearance of maternal mRNAs or transcripts from transposons.

### Enrichment of transposable elements (TEs) in promoters

We systematically studied the distribution of all repetitive elements (REs) in the mouse genome based on RepeatMasker data available at UCSC genome browser website^51^. Covering 44% of the genome, the 5,147,736 REs are classified into 1554 different types with unique names and consensus sequences in the RepBase database^54^. They belong to 47 repeat families which are grouped into 16 repeat classes.

Figure 2A shows highly enriched REs in the 2kb promoters of 10 gene clusters. B1 and B2 SINE elements are overrepresented in the promoters of Cluster C, D, F and G genes. Most of these genes are upregulated at the 2C stage (Figure 1B). This is congruent with the fact that transcription of B1 increases at the 2C stage. For example, 32.7% of genes in cluster C contain at least one B1_Mus2 element in their promoter, which is much higher than the percentage (15.7%) in cluster A. Note that B1_Mus2 is just one of many forms of B1 elements.

**Figure 2.**
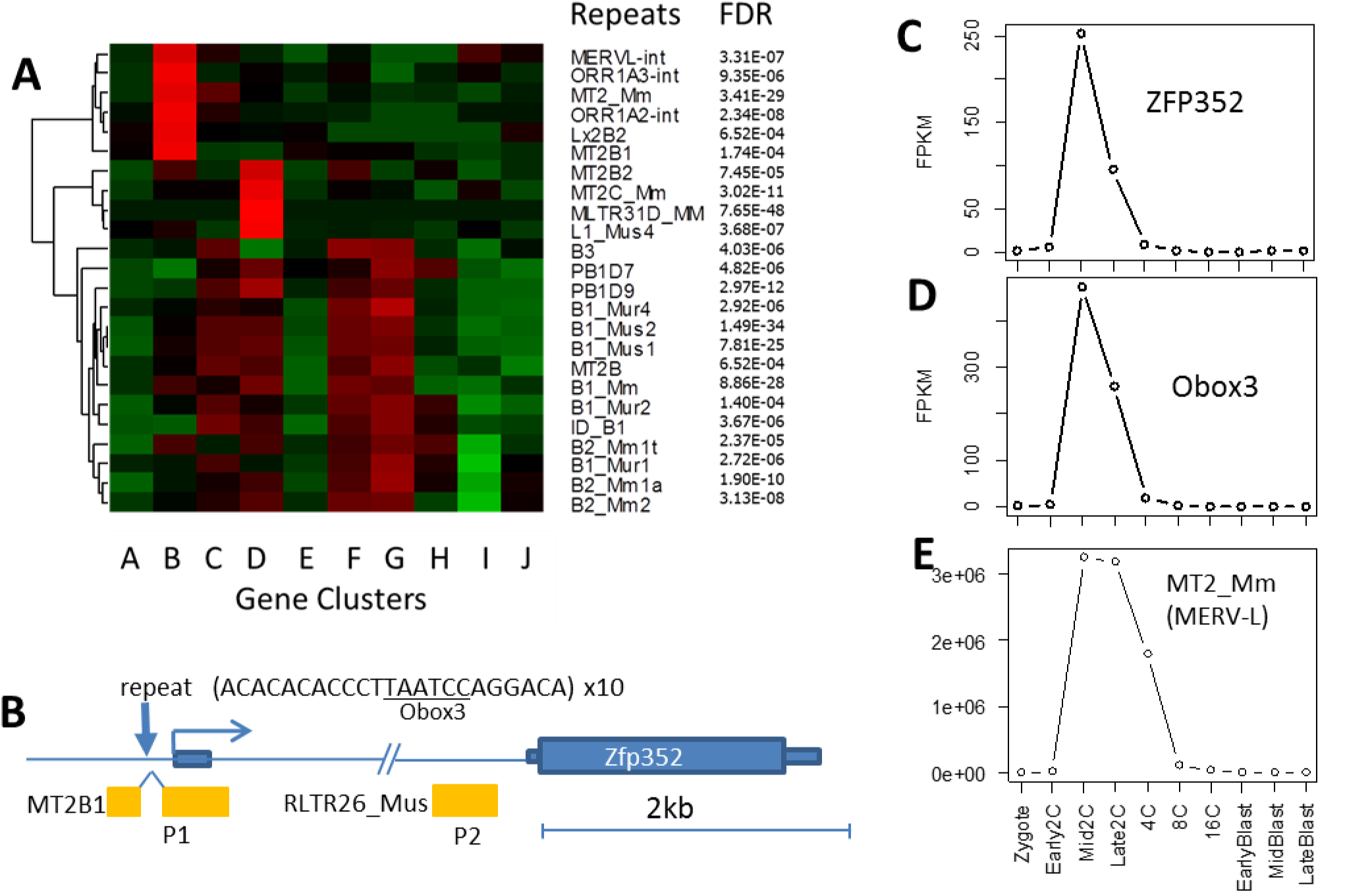
A) Enrichment of repeat elements in the promoters of genes. Original data matrix represents percentage of genes in a cluster containing a repeat in their promoter. Red indicates a certain repeat is enriched in the promoters of the gene clusters. B) *ZFP352* gene contains a LTR element (MT2B1) around the translation starting site. Another LTR element (RLTR26) is in the intron. These two positions correspond to known promoter regions marked as P1 and P2. Expression levels of Zfp352, Obox3, and retroelement MT2_Mm are shown in C, D and E, respectively.

Promoters of Cluster D genes are 9.6-fold enriched with MT2C_Mm LTR compared with other genes (false discovery rate^55^ of 3.02 × 10^-11^). The MT2C_Mm retrotransposon itself is transcribed between 2C and 16C, similar to Cluster D genes. MT2B2 and MLTR31D are also overrepresented in the promoters of cluster D genes. Although relatively enriched, these LTRs are only found near a small proportion of Group D genes. As shown in Supplementary Table S5, about 5% of Cluster D genes contain MT2C_Mm elements, which is a 6.7-fold enrichment compared to all other clusters combined.

Promoters of Cluster B genes are enriched with other LTR elements, namely MERVL-int, ORR1A2-INT, ORR1A3-INT, MT2_Mm, and MT2B1. The most significant is a 9.6-fold enrichment of MERV-L (MT2_Mm) with FDR < 3.41 × 10^−29^, compared with genes in other groups combined. This retrotransposon itself is transcribed at very high levels only in 2C-4C stages (Figure 2E), similar to Cluster B genes, suggesting a shared mechanism of regulation. This agrees with previous reports^32^. Supplementary Table S2 includes 117 genes that contain ERVL elements in promoters and show an expression pattern similar to these LTRs.

As an example, transcription of *Zfp352* (zinc finger protein 352) starts at the middle of the LTR element MT2B1 (Figure 2B). This gene is sharply upregulated at mid 2C stage, before quickly decreasing to very low levels at the 4C stage (Figure 2C). This gene has been studied experimentally by Liu *et al.*^56^, who found one major promoter P1 and an alternative, weaker promoter P2 in the intron. P1 actually lies within the MT2B1 element and P2 is in another LTR in the intron (Figure 2B). *Zfp352* is likely generated by retrotransposition, similar to homologous pseudogene Zfp353-ps^57^, where mRNA sequences are reverse-transcribed and inserted into the genome. Retrocopies of mRNA typically have no intron and are not expressed due to the lack of a functional promoter. But a subsequent or preceding insertion of LTR elements upstream of the retrocopy can provide a promoter. The expression of 117 such genes (See supplementary Figure S8 for another example, Gm9125) was observed during PD when LTRs are actively transcribed. These expressed retrogenes can be further studied.

### Early genes share motifs for Obox homeobox factors

We scanned the proximal promoter region (-300, 50bp) of all the genes for transcription factor binding sites (TFBS) using the comprehensive CIS-BP database^58^, which includes thousands of binding motifs determined using protein binding microarrays or inferred based on shared protein domains. Out of the 1823 binding motifs scanned, four were found enriched in the most highly induced gene group at mid 2C (Figure 3A). Sharing the same “TAATC” core sequence, these four motifs can be bound by OBOX1, OBOX3, OTX1, PITX2, and other factors. Noticeably, the *Obox3* gene is upregulated by more than 100-fold at mid 2C, and quickly downregulated at 4C (Figure 2D). This is similar to the expression pattern of its potential target genes such as *Zfp352* (Figure 2C), suggesting a potential regulatory role of Obox factors.

**Figure 3.**
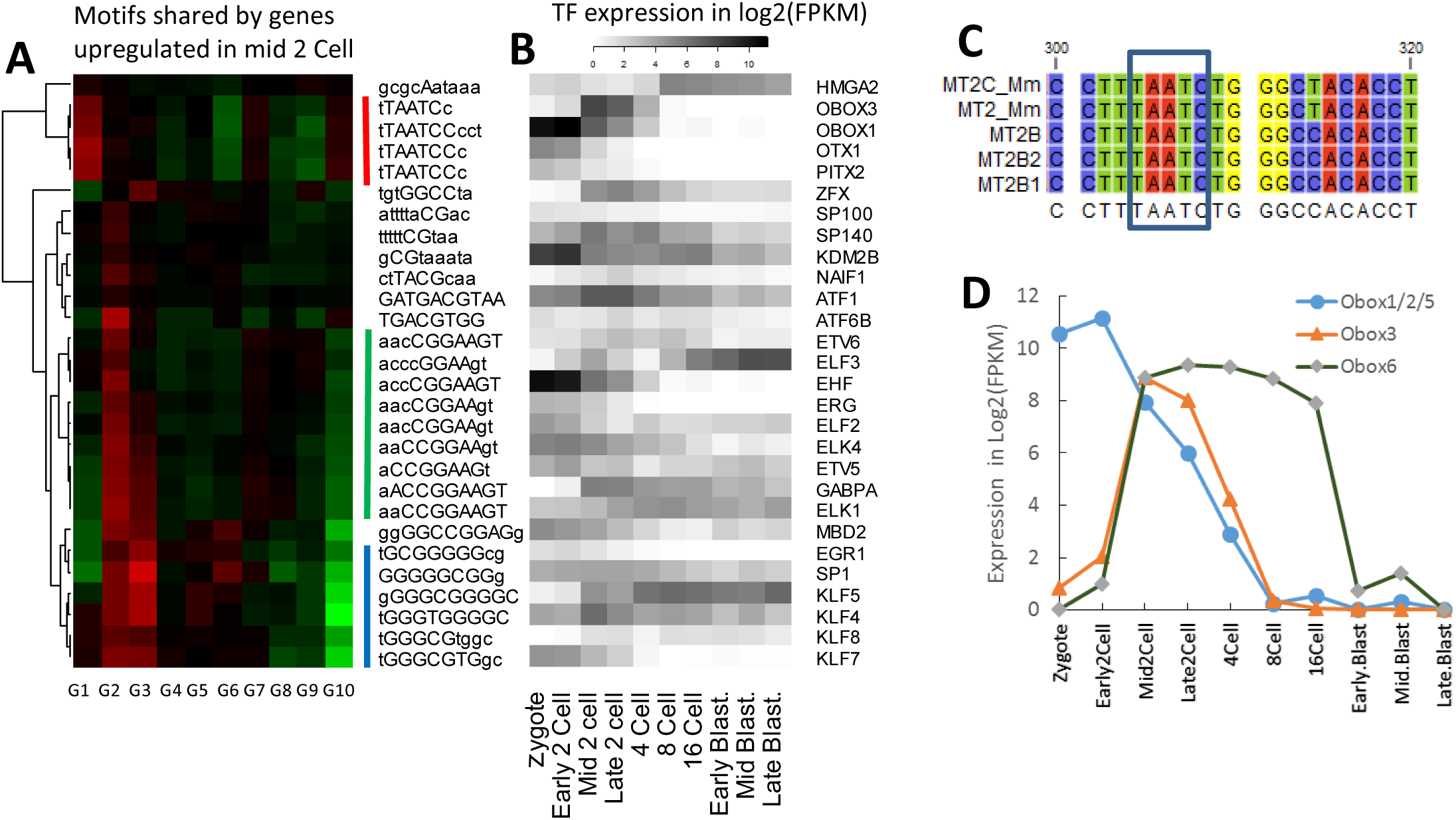
Enrichment of transcription factor binding motifs in genes highly induced at mid 2-cell stage. A. Enriched motifs in the promoters of genes with highest fold-change at mid 2C. G1-G10 represents 10 groups of 500 genes sorted by fold-change. Red indicates enrichment of motifs. B) The expression patterns of TFs that bind to the corresponding motifs. C) Binding motifs by Obox families in LTRs belong to the ERVL family. D. Expression patterns of Obox family homeobox factors are highly regulated during PD.

Oocyte specific homeobox (Obox) family transcription factors (TFs) have not been studied in detail because they seem to be rodent-specific and are only expressed during PD^59^. They are highly regulated in oocytes and cleavage stage embryos (Figure 3D). They are among the top 10 TFs when ranked by variance in gene expressions during PD. Since these TFs bind to similar motifs (supplementary Figure S7), it is difficult to distinguish the true driver of gene expression. Three members (Obox1/2/5) are similar in sequence, and are treated as one group in our RNA-seq mapping; they are highly expressed in oocytes and are maternally derived. Due to their higher expression at 2C and shared binding motifs, Obox1/2/5 could work together with Obox3 to regulate gene expression. Experimental studies are needed to confirm whether these factors are redundant or have differences. Obox6 expression is elevated from 2C to 16C, and low in the blastocyst stage (Figure 3D), in agreement with a previous study^60^.

We also examined the expression of these genes in another single-cell RNA-seq dataset^13^, and found that the patterns are similar, except that Obox3 is expressed lower (See supplementary Figure S51). Using this this dataset, we also identified the enrichment of the same “TAATC” motif among genes upregulated in 2C stage (See supplementary Figure S52).

Obox binding sites are enriched in ERVL family LTR elements that are overrepresented in the promoters of cluster B genes (Figure 3C). For example, the 493 bp consensus sequence of MERV-L contains three Obox binding sites, while the 521 bp MT2B1 contains four. Although not as highly transcribed as MERV-L, MT2B1 is one of reliable predictors of nearby gene expression among LTRs, as the corresponding transcripts almost always start right from the repeat, similar to *Zfp352*. Interestingly, the MT2B1 element in the main promoter of *Zfp352* is interrupted by a (10x) tandem repeat of 21 bp sequence, which contains the Obox binding motif (Figure 2B). These tandem repeats may be selectively retained during evolution. Furthermore, genes with multiple Obox binding motifs are more highly induced at mid 2C when compared to genes with one or no such motifs (supplementary Figure S10). The retrovirus-like LTRs contain their own promoters and enhancers. These promoters may be bound by Obox TF families to drive the expression of both retrotransposons and nearby genes.

Among the genes with transient 2C expression is the Zscan4 family (Zscan4b, Zscan4c, Zscan4d, Zscan4e, and Zscan4f). Zscan4 proteins are involved in telomere elongation and genomic stability in ESCs^61^, and have been reported to restore developmental potency in ESCs^62^ and to transiently activate embryonic genes in induced pluripotent cells (iPSCs)^63^. Among the genes transiently expressed at 2C, Zscan4 genes ranked highest in terms of number of Obox3 binding sites, with three binding sites at 90% similarity level in the 350 bp around the TSS. Further study is needed to verify if Obox family TFs are upstream regulators of Zscan4 genes, and whether induction of Obox gene expression in adult cells could promote pluripotency.

Thus, there is evidence that the poorly-characterized Obox family induces gene expression at mid 2C to jump start ZGA. Even though Obox6 mutants develop normally^60^, loss of function of other family members may be lethal to the embryo, due to potential effects on hundreds of downstream genes and even LTR transposons.

The overrepresented “TAATC” motif can also bound by other homeobox TFs like OTX1 and PITX2 (Figure 3B and C). These TFs may also regulate gene expression, as their transcripts are also present in oocyte, even though at a much lower level (Figure 3B). Recently, a similar motif was discovered by Töhönen *et al*. ^15^ in the promoters of 32 human genes upregulated in early ZGA using single-cell RNA-seq. This agreement may suggest a conserved mechanism of ZGA, and warrants experimental confirmation.

### SINEs associated with broad genome activation

To further study expression change in late 2C, we ranked genes by their fold-changes from late 2C compared to mid 2C, and then divided them into 24 groups of 500 genes. As shown in Figure 4A-C, the REs differentially distributed in the promoters of these gene groups are mainly SINE elements, including the Alu family (mainly B1) and B2 family. The most significantly associated elements are B1_Mm, B1_Mus1, and B1_Mus2 (Figure 4C). These B1 elements are highly similar in their sequences, with only a few base-pair differences in most cases (supplementary Figure S11). With over 400,000 copies, B1 is one of the most prevalent retrotransposons in the mouse genome. Similar to human Alu elements, B1 originated from initial duplication of the 7SL RNA^64^. The distribution of SINE, but not LINE, is conserved across species^65^. B2 elements originated from tRNAs^64^. It is possible that B1 and B2 elements play a role in gene regulation during PD, and their fast expansion may have benefited mammalian development.

**Figure 4.**
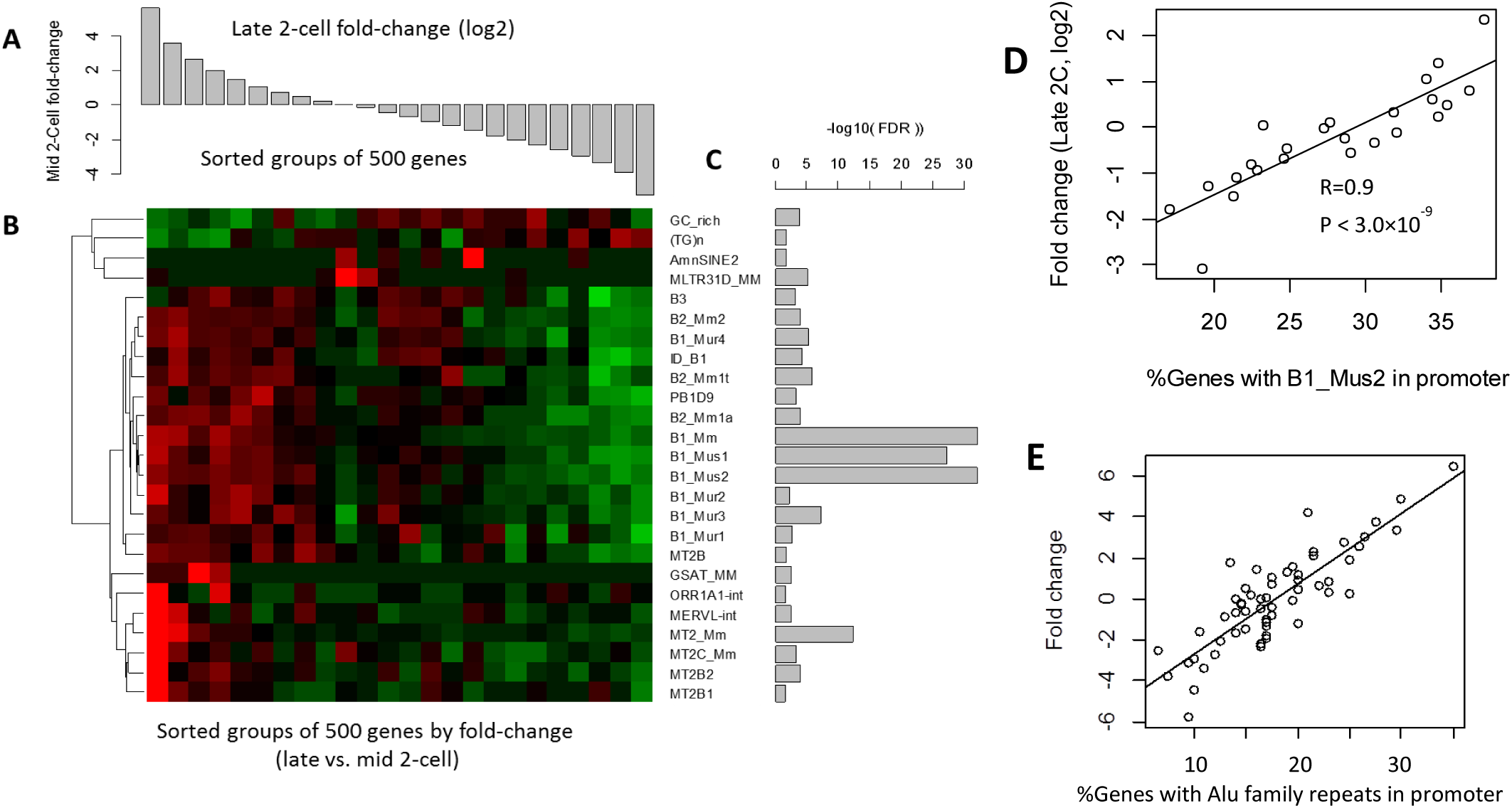
Enrichment of SINE elements in genes upregulated in the late 2-cell stage. A) Average fold-change for 24 groups of 500 genes. B) Relative enrichment (red) or depletion (green) of repeat elements in the promoters of genes in these groups. C) FDR values derived from Chi-square tests of independency of repeat element frequency and gene groups. D) Correlation of fold-change in 2C with the presence of B1 elements in gene groups. Each point in the plot represents one group of 500 genes. The vertical axis represents the percentage of genes in the group with B1_Mm in their promoters, while the horizontal axis represents average fold-change. E) Association of repeats with fold-change during the late 2-cell stage. The average fold change of genes with one or more repeats within 2kb on either sense or antisense strand.

Figure 4B also shows that genes downregulated at the 2C stage are depleted in these elements. The average fold-change observed in these groups is highly associated with the percentage of genes containing Alu family elements in their promoters. As shown in Figure 4D, the Pearson’s correlation coefficient (PCC) is 0.90 (P < 3.0×10^−9^) for one of such element B1_Mm. About 36% of the 1500 genes that are upregulated by more than 2 fold contain B1_Mm elements, which is much higher than the 18% observed in gene downregulated by 2 fold. In addition to B1_Mm elements, other B1 and B2 elements also show this trend. Using a different dataset of single-cell RNA-seq data^13^, we were able to confirm this remarkably linear and consistent correlation (Figure 4E).

Multiple B1 elements in promoters are associated with stronger upregulation in a dosage-dependent manner (Figure 5A-D). As shown in Figure 5C, the 496 genes with five B1 elements or more (some are partial) in promoters are upregulated by 2.1-fold on average, which is significantly higher than the 1.65-fold observed in 568 genes with 4 B1 elements (P<0.0064). This is in turn higher than genes with three B1 elements (P<0.027). The effect of B1 elements is surprisingly linear. Each additional B1 element is associated with an approximately 20%-40% upregulation (See supplementary Table S1). Among genes that are upregulated more than 2 fold at late 2C stage, 59% contain Alu family repeats in promoter, which is much higher than the 41% observed other genes.

**Figure 5.**
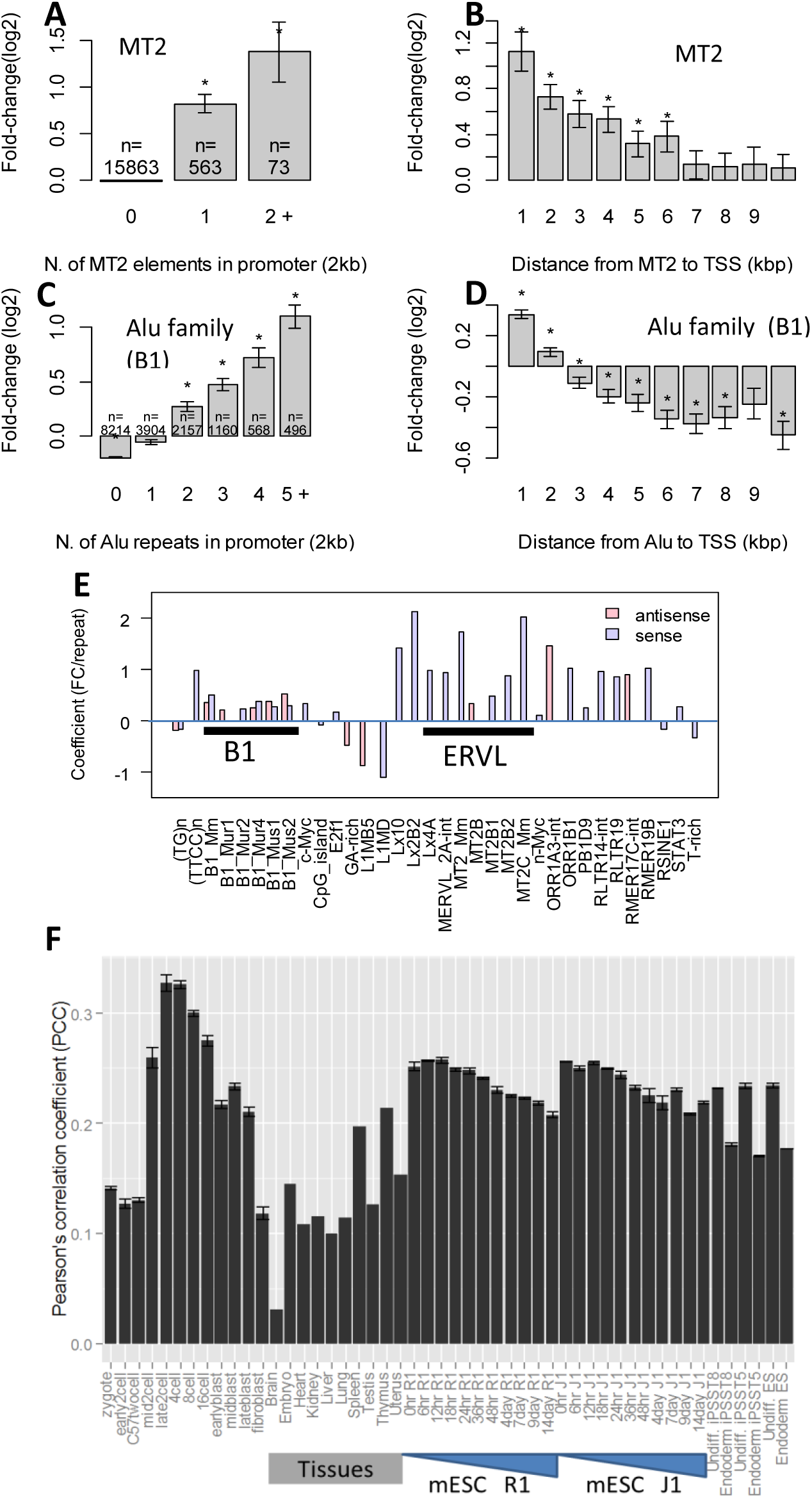
Presence of some repeats strongly correlated with activation of gene expression during 2C stage in a dosage- and distance-dependent manner. The heights of the bars indicate average fold-change in late 2-cell stage compared to early 2-cell. Significant deviations from zero are indicated by stars. The numbers on the bar represent the number of genes affected. The MT2 repeats in A) and B) can be MT2B, MT2B1, MT2B2, MT2_Mm, or MT2C_Mm. In C) and D), Alu family repeats mostly represent B1 elements. E) Correlation of the number of Alu family repeats in promoters and broad gene expression in various cells and tissues. Error bar shows standard error calculated from replicates.

The effects of the B1 elements are also dependent on the distance to TSS (Figure 5D). The correlation is stronger when B1 elements are located closer to TSS. Further upstream, the effect of Alu elements is weaker, diminishing after 5-7kb. This is similar to what was observed in the MT2 elements (Figure 5B). B1_Mm and other mouse-specific B1 elements are more efficient than other Alu family repeats common in muridae (B1_Mur1-4) or rodentia (PB1D9). But it is difficult to isolate a specific type of element, as different subtypes of Alu family repeats often co-appear.

To systematically investigate the effects of various repeats on ZGA, we used multiple linear regression to model 2C fold change as a function of the numbers of various kinds of repeats within the 2kb promoter. The model also included CpG islands and TFBS of 13 TFs determined by Chip-Seq^66^. The results (Figure 5E) confirm the effect of SINE and LTR elements. While the effects of ERVL family LTRs are strong and only seen on the same strand, B1 elements have weaker effects on both strands. Genes with multiple c-Myc sites show a bigger fold-change. E2F1 binding sites are associated with weaker but significant upregulation in a larger number of genes. The effect of Alu elements and c-Myc and E2F1 binding sites can be confirmed using independent single-cell RNA-seq data^13^ (supplementary Figure S12).

SINEs are a major source of CpG dinucleotides in mammalian genomes. It is possible that the effect of Alu elements in promoters is through the contribution of CpG sites that affect epigenetic modifications. Linear regression analysis shows B1 elements have a much more significant (P < 2.2×10^-16^) correlation with ZGA fold-change than those of CpG dinucleotides (P < 0.04), or CpG islands (P<0.004). Also, gene clusters defined by methylation of promoter regions during PD^19^ (supplementary Figure S18) have little in common with the gene clusters by expression (Figure 1B). Therefore, the effect of B1 repeats cannot be fully explained by CpG sites.

#### SINEs correlate with gene expression in adult tissues and stem cells

It has been reported that Alu family repeats are enriched near housekeeping genes^67,68^, which are both broadly and highly expressed across tissue types. We calculated the correlation coefficient between expression levels and the number of Alu family repeats in promoters among genes. As shown in Figure 5E, there is a significant positive correlation for all cells and tissues. The PCC dramatically increases from 0.13 at early 2C to 0.33 at late 2C, and gradually decreases to 0.21 at the late blastocyst stage. PCC is higher in embryonic cells than in adult tissues. This association is independent of CpG islands, which also occur in association with higher gene expression (supplementary Figure S19).

Occurrences of Alu family repeats are a stronger predictor of gene expression in undifferentiated iPSCs compared with day 5 definitive endoderm^69^. A similar pattern of correlation is observed with B2 family repeats (supplementary Figure S20). SINEs may play a role in regulating gene expression in both pre-implantation embryos and ESCs.

Urrutia *et al.*^70^ found no association between Alu content and peak (maximum) gene expression level across tissues. However, we found a significant correlation (R=0.16, P < 2.2×10^-16^) with average gene expression. In addition, even among genes expressed in all tissues, more Alu elements in the promoter are associated with higher average expression (R=0.13, P < 2.2×10^-16^). The same is true for tissue-specific genes (R=0.16, P < 2.2×10^-16^).

#### Promoters of mESC–specific genes are rich in SINEs and low in CpG_island

To further delineate the role of Alu family repeats in gene regulation, we re-analyzed RNA-seq data^71^ of normal tissues from both fetal and adult mice, as well as embryonic stem cells (mESCs). We divided 16,989 protein-coding genes into 25 groups using k-means clustering based on their expression pattern in various tissues/cells (Figure 6A). In addition to ubiquitously-expressed housekeeping genes, we identified many clusters of tissue-specific genes. Cluster 13 contains 89 genes specifically expressed in mESC cells. As shown in supplementary figure S50, this includes known TFs (Nanog and Pou5f1), as well as many other factors such as Zscan10, Esrrb Foxn4, and Sox15. These genes contain many Alu family repeats in their promoters (-5kbp to 1kb). As shown in Figure 6B, their average Alu/B1 coverages are as high as housekeeping genes, much higher than other tissue-specific gene clusters. Unlike the promoters of housekeeping genes, the promoters of mESC-specific genes are less likely to have CpG islands. For example, the Alu-rich promoter regions of Nanog and Oct4 are shown in Figure 6C. Human orthologs of these two genes are also enriched with Alu elements. SINE elements might be important for the expression of pluripotency related genes.

**Figure 6.**
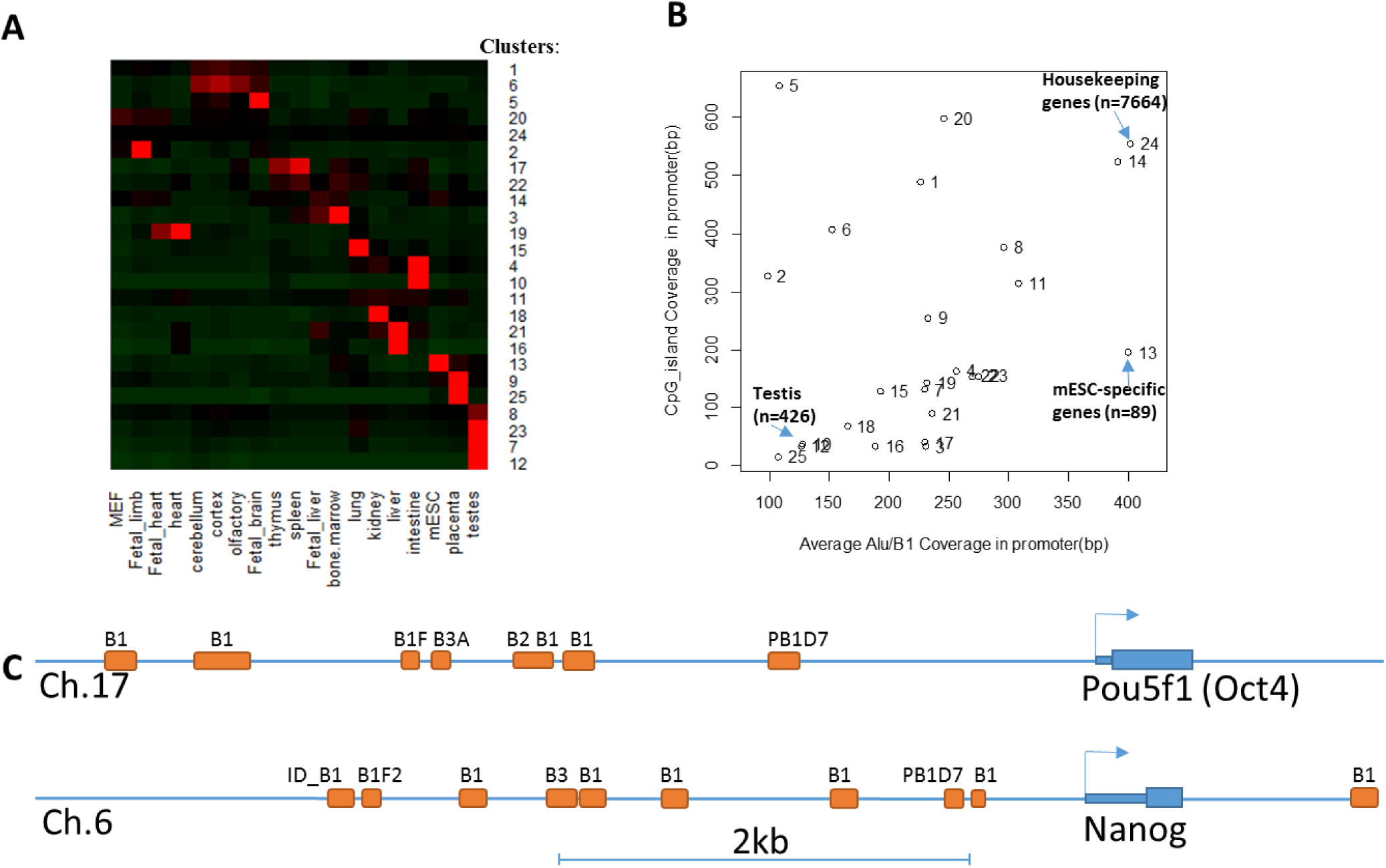
Genomic content of promoters of mESC-specific genes. A) Results of k-means clustering of normal tissue gene expression show housekeeping genes (Cluster 24) and tissue-specific genes. B) Gene clusters are plotted by average coverage of Alu/B1 repeats and CpG island coverage. Housekeeping genes (Cluster 24) are high in both Alu/B1 coverage and CpG island. mESC specific genes (Cluster 13) are high in Alu/B1, but low in CpG island. Gene Clusters specifically expressed in testis, intestine, and placenta are lower in both. C) Devoid of CpG islands, the promoters of Pou5f1 and Nanog are enriched with Alu/B1 elements.

### SINEs correlate with ZGA in other species

Single-cell RNA-seq data of early embryogenesis in humans^72,73^ were used to investigate the correlation between transposons and ZGA, which occurs between 4-and 8-cell stages^74^. Regression analysis at the repeat family level shows a significant association between the presence of Alu family repeats (FDR = 1.74×10^-50^) and gene expression (Table S2 in supplementary document). The most prevalent Alu element, AluJb, is significant on both sense and antisense strands. Presence of an Alu element is associated with 15-48% upregulation. Figure 7A shows that genes with multiple Alu elements in promoters are upregulated in a dosage-dependent manner, similar to what was observed in mouse. As suggested by Figure 7A-B, genes with fewer than two Alu elements in the promoter are downregulated at 8C, similar to what was observed in mouse. This is in agreement with another single-cell RNA-seq analysis of human preimplantation embryos, which shows that Alu elements are enriched in upstream of TSS of the 129 genes upregulated during PD^15^. Alu elements were found to contain binding motifs for PITX1 and TBX1^15^.

**Figure 7.**
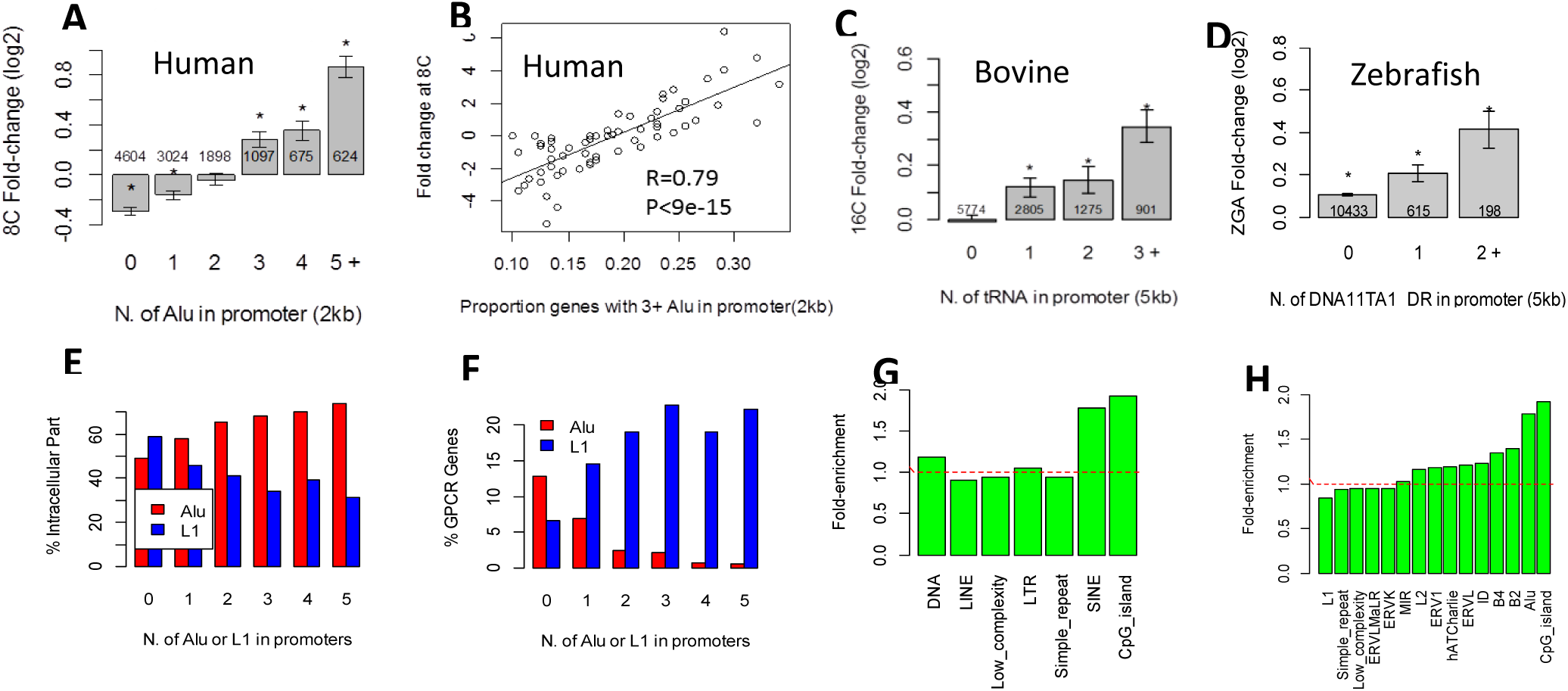
A) Genes with more Alus in promoter are upregulated in a dosage-dependent manner. Stars indicate significant difference from zero. B) Among gene groups defined by fold-change, highly expressed ones tend to contain more Alus in promoters in humans. C) Association of gene expression with tRNA family of SINE repeats in the bovine genome. Stars indicate significant difference from zero. D) DNA transposon, DNA11TA1_DR, is associated with gene upregulation during ZGA in zebrafish in a dosage-dependent manner. E) Mouse genes with multiple Alu family repeats, mostly B1 elements, are associated with GO:0044224, intracellular part. Genes with L1 elements in their promoter, on the other hand, are depleted in genes related to intracellular part. F) Genes with L1 elements are enriched with GPCR activity, while in genes containing Alu elements, such genes are depleted. G) Mouse genes with SINE element in promoter are enriched in genes with yeast orthologs (>20% identity according to BioMart). H) Among SINE elements, Alu family, mostly B1 elements, are enriched for genes with yeast orthologs.

We also compared the occurrences of human Alu and mouse B1 elements in promoters of orthologous gene pairs. The number of Alu elements in human gene promoters is highly correlated with the number of B1 elements in orthologous mouse gene promoters (R=0.57). This has been noted as surprising, as B1 and primate Alu elements replicated in these genomes independently^75^. The differences in ZGA fold-changes between orthologous gene pairs in human and mouse are significantly associated with the differences in the number of Alu family repeats in their promoters (P < 1×10^-14^). The rapid expansion of B1 in the mouse and Alu in the human genome may contribute to gene expression divergence.

Bovine ZGA is associated with tRNA SINE repeats. We used RNA-seq data based on pooled embryos^12^. The repeat families associated with expression change during ZGA between 8C and 16C are ERV1, simple repeat, and tRNA. The tRNA family SINE repeats in bovine have a weaker but significant association (FDR< 1.1×10^-5^). It is also dosage-dependent (Figure 7C), as genes with three or more tRNA family repeats are more highly upregulated than genes with one or two such repeats (P<0.011). The tRNA family repeats are associated with higher gene expression in both sense and antisense strands. The most significant repeats are SINE2-1_BT and SINE2-2_BT, which are 120bp bovine-specific SINE repeats derived from tRNA. The association with tRNA family repeats can also be confirmed using DNA microarray data^9^.

ZGA in zebrafish is associated with AT-rich DNA transposons (supplementary Figure S23). The zebrafish (*Danio rerio*) genome^76^ is dominated by more than 2 million DNA transposons. There are fewer retrotransposons compared with mammals. Using RNA-seq, Harvey *et al.* studied the zebrafish ZGA^77^, which happens at about 3.5 hours post-fertilization. Results from regression analysis (Tables S3 and S4 in supplementary documents) show that some DNA transposons are highly associated with gene upregulation at ZGA. The most significant is DNA11TA1_DR, a non-autonomous DNA transposon. There are 813 genes containing this repeat in the 5kb promoter region, and their expression is significantly higher than other genes (FDR<1.88×10^-9^). The 198 genes with two or more DNA11TA1_DR elements are induced at significantly (P<0.03, t-test) higher levels than the 615 genes with one element, which is in turn higher (P<0.013) than genes without such an element (Figure 7D).

The accumulation of SINEs near genes is likely to result from a positive selection process^35^. Since there is no known mechanism to remove SINEs from genomes, it is difficult to explain the lack of SINEs in gene-poor regions^35^. The transposons significantly correlated with ZGA are often prevalent in and specific to the host species. Specific transposons seem to be encouraged to expand during the course of evolution.

### Non-random distribution of transposons in the genome

We studied the distribution of all repeats across the mouse genome using the EBI approach. While some REs like B1 are prevalent, others are only observed dozens of times. We found that the frequency follows lognormal distribution (see supplementary document). Lognormal distribution implies that growth rate is independent of existing occurrence^78^. The distribution of the distances between repeats is power-law like^79^, which could be expected as transposons often “copy-and-paste” to nearby loci and form clusters on the genome.

Some repeats show enrichment and strand-preference near genes (supplementary Figure S29). We found that SINEs are enriched in introns, promoters, and downstream regions, suggesting that SINEs are located near genes. On the contrary, LINEs are depleted from these regions and are away from genes. There are 97 types of LTRs that are specifically enriched in promoter regions. Some repeats demonstrate strand-preference. For example, RLTR10-int repeats align 4.6-times more frequently on the same strand relative to the nearby gene than on the opposite strand, which is highly unusual (P<1.4×10^-74^). There are a total of 14 repeats that are enriched in a strand-specific manner, including several prevalent ERVL-MaLR family members (MTC, ORR1D1, ORR1A2, ORR1A2-int), ERVK family members (RMER19B, MYSERV6-int, MYSERV-int, RLTR10, RLTR10-int, MLTR18A_MM, and RLTR9A3A), the ERVL family (MT2B1, RMER15-int), and the ERV1 family (LTR72_RN). This could be explained by new LTR elements generating new genes by activating retrogenes, as discussed. We have shown that MT2B1 contains Obox3 binding sites and is strongly associated with gene expression during PD. Other elements may regulate gene expression in other situations^80^.

Surprisingly, *most intronic retrotransposons are more likely found on the opposite strand of the host gene* (supplementary Figure S29). It is possible that intronic sequences, once spliced off transcripts in the nucleus, are spliced, reverse-transcribed, and inserted back into the genome, resulting in intronic retrotransposons on the antisense strand. DNA transposons do not show such a strong strand-specificity. More investigation on the intronic strand-specificity is needed to verify this possible mechanism.

#### Genes with multiple B1 in promoters form core cellular machinery

Based on Gene Ontology (GO)^81^, we found that genes with multiple B1 elements in promoters are more likely to code for proteins that constitute intracellular parts (Figure 7E) and less likely to be related to G-protein coupled receptor (GPCR) activity, an extracellular signaling process (Figure 7F). On the contrary, genes with L1 LINE elements in their promoter are enriched in GPCR related genes and depleted of intracellular parts. Low complexity repeats are found to be enriched in promoters of genes related to the RNA metabolic process (supplementary Figure S30). Thus, the distribution of repeats in the genome is related to the function of genes. It is possible that the mouse embryo utilizes B1 elements to quickly establish core proteins during ZGA.

Mouse genes with SINEs in promoters are more evolutionarily conserved. Figure 7G-H suggests that mouse genes with SINEs in the promoter are much more likely to have yeast orthologs (>20% identity according to BioMart^82^). This is similar to, but independent of, the effect of CpG islands (supplementary Figure 30). A similar trend is observed in human genes (supplementary Figure S31). Thus, the distribution of transposons is correlated with the function of nearby genes. The distribution of transposons may be regulated through selection.

### Key transcription factors in early development

Taking advantage of high-resolution expression data^14^, we also systematically analyzed TFBS in the promoters of genes co-regulated at other stages beyond 2C. Enriched TFBS and expression patterns of corresponding genes are shown in supplementary Figures S32—S49, which are summarized in Figure 8A.

**Figure 8.**
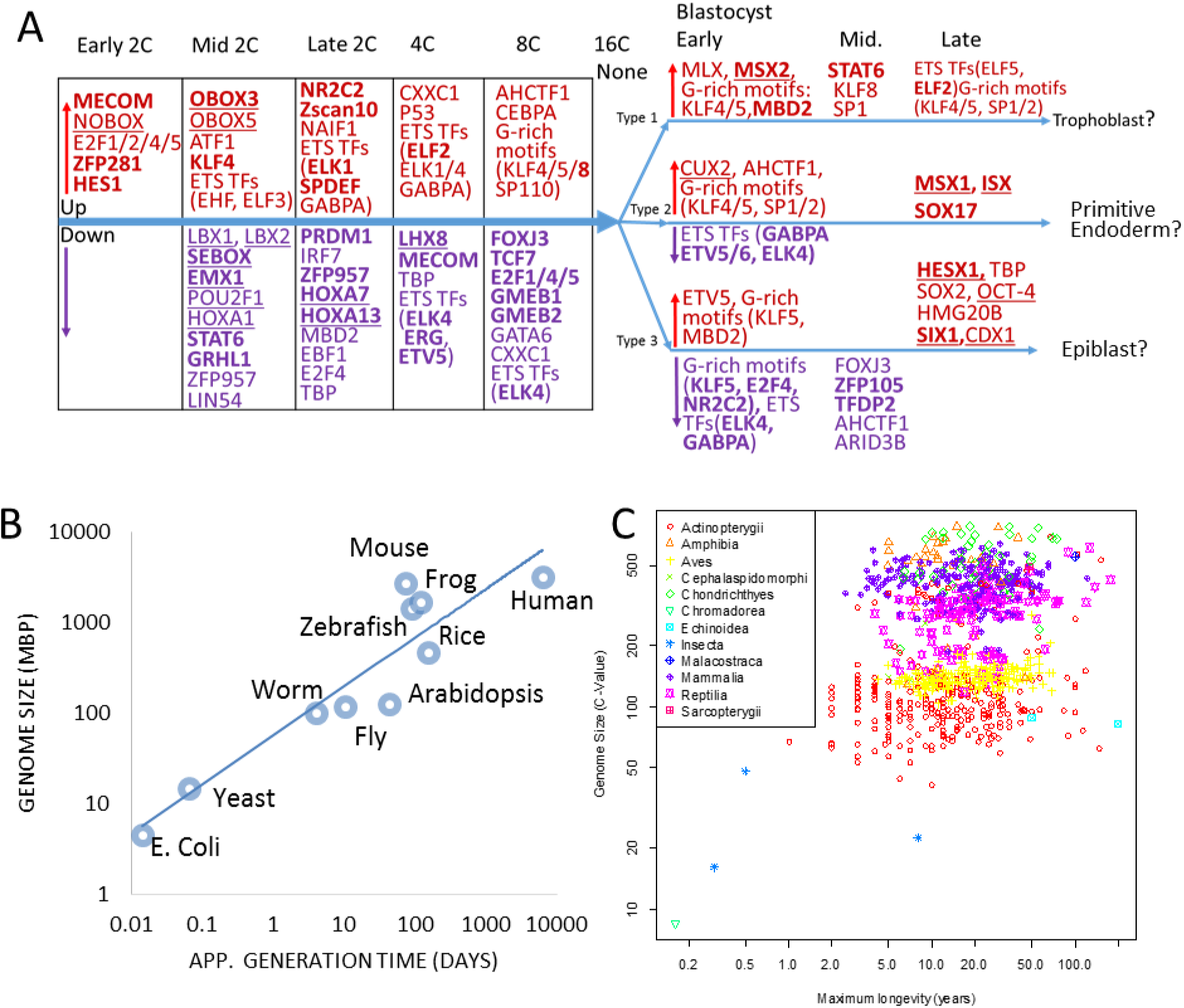
A) Transcription factors (TFs) identified during PD. Upregulated and downregulated factors are shown in red and purple, respectively. Homeobox TFs are underlined. Genes coding for the TFs that are upregulated or downregulated at the same stages provide more confidence and are shown in bold. B) A possible correlation between genome size and generation time across selected model organisms. C) A possible correlation between genome size and longevity among animals. Both axises are in log scale. Taxonomical classes are represented by different shapes and colors. N = 939, R = 0.245, P=2.4×10^−14^.

We identified several TFs known to regulate embryo development, such as SOX2, OCT4 (POU5F1), and KLF4^83,84^. In addition, many TFs in Figure 8A are upregulated at the same developmental stage as their potential target genes, thus giving more support to their involvement in gene regulation. This includes Obox3 and KLF4, which are upregulated at mid 2C, as well as NR2C2, Zscan10, and ELK1 at late 2C. Zscan10 is known to be expressed during PD^85^ and is involved in maintaining pluripotency in ESCs^86^. Similarly, many motifs enriched in the promoters of downregulated genes are bound by TFs that are downregulated at the same time. Some of the TFs are likely maternally derived with high expression in the zygote and reduced at the 2C stage: LBX1, LBX2, SEBOX, ZFP959, POU2F1, STAT6, GRHL1, EMX1, PRDM1, HOXA7, LHX8, EBF1, and E2F4. SEBOX is known as one of the maternal effect genes (MEGs), and its RNA products are carried over from oocyte to regulation of gene expression in early PD^87^. Further study should verify whether other TFs in this list are MEGs.

The GGAA motif bound by the E26 transformation-specific (ETS) domain TFs is repeatedly identified as enriched at several stages. One of the TFs, GABPA, has been shown to be involved in early embryogenesis^88^. The most highly expressed ETS domain TFs in oocyte and 2C is EHF. ELF3 is highly expressed at the blastocyst stage. ETS domain TFs are a large and conserved family of TFs involved in a variety of developmental processes in animals^89,90^.

Similarly, the G-rich motifs are repeatedly identified at various stages. This motif can be bound by KLF4 or other TFs, such as KLF5, KLF8, SP1, SP2, ZBTB7B, etc. The most highly expressed is KLF5, which was reported to regulate lineage formation in the pre-implantation embryo^91^, alongside KLF4^92^.

To account for the heterogeneity among cells in the blastocyst stage, we used hierarchical clustering to divide them into three types/lineages, which were then analyzed separately. Sox2 and OCT4 binding sites are enriched in promoters of genes upregulated in Type 3 cells at the late blastocyst stage. These markers indicate that type 3 cells likely correspond to the epiblast, which is derived from the inner cell mass (ICM) and leads to the embryo proper^93^. Figure 8A shows that two homeobox TFs, HESX1, and SIX1, are both upregulated together with their target genes in this type of cell. HESX1 is believed to be downstream of multiple pluripotency related pathways^94^. SIX1 is considered to be an oncogene and is known to be involved in embryonic muscle formation^95,96^.

In Type 2 cells, SOX17 binding sites are overrepresented in promoters of genes upregulated at the late blastocyst stage, when the SOX17 gene itself is also upregulated. SOX17 directly promotes differentiation towards extraembryonic cells, which leads to the primitive endoderm^97^, which is also derived from ICM but develops into the yolk sac. Therefore, type 2 cells are likely committed to the primitive endoderm. MSX1 and ISX are two other homeobox TFs identified as inducing gene expression in these cells. Little is known about ISX in embryogenesis, but it is highly induced at the late blastocyst stage in type 2 cells. Interestingly, we found *MSX1* and *MSX2* are associated with primitive endoderm and trophoblasts, respectively. Their differential role should be further studied.

Type 1 cells may represent trophoblasts, which make up the outer layer of the blastocyst. In the promoter of genes highly induced in this type of cell, we failed to detect binding sites for CDX2, a key regulator for trophoblasts ^93,98^. But sites bound by MSX2 are enriched. MSX1 and MSX2 were shown to be critical for the interaction between the blastocyst and the uterus^99,100^. These two proteins are highly conserved in mammals^101^, and mutations of *Msx1* and *Msx2* lead to failures in implantation^99,100^.

Our analysis identified many other TFs that could potentially contribute to the complex gene regulatory network during PD. For example, *MECOM* is upregulated at early 2C and downregulated at 4C, along with its potential target genes. MECOM is highly expressed in the embryo, and mutant is embryonic lethal^102,103^. Figure 8A also shows that we identified many homeobox TFs, which are believed to be regulators of morphogenesis and development^104^. In addition, many TFs in Figure 8A have been studied in relation to embryogenesis and cancer. Further study of these TFs will elucidate their role in gene regulation in PD.

## Discussion

### Transposons, mutagenesis, and the C-value paradox

Some organisms in our ecosystem can finish a reproduction cycle in 20 minutes, while others require more than 10 years to reach sexual maturity. If genetic mutations happen stochastically at similar rates, a huge imbalance in how quickly organisms evolve and adapt would result. Slow-reproducing organisms, therefore, are under pressure to find ways to dramatically promote mutagenesis and genotype diversity. Transposition of mobile DNA elements may be a necessary “copy-and-paste” mechanism that promotes not only insertional mutations but also homologous recombination. We showed here that they may also regulate many coding genes during early development and thus will have substantial influence of morphogenesis. Some retrotransposons are even active in somatic cells and lead to genetic mosaicism within individuals^105^. There is additional evidence that TEs are drivers of genome evolution^32,34,46,106^, rather than just “junk” DNA.

Following this argument, we would expect slow-reproducing organisms to have more TEs in their genome, leading to larger genomes. Across model organisms, generation time seems to be proportional to genome size on a log-scale (Figure 8B). Using larger datasets^107,108^, longevity and genome size are weakly but significantly correlated (PCC = 0.245, P=2.4×10^−14^) across 939 animal species (Figure 8C). Further study is needed to confirm the correlation in more organisms. But this may shed some light on the C-value (genome size) paradox^109,110^: eukaryote haploid DNA contents vary greatly, but are unrelated to organismic complexity. Even though TEs can be disruptive for individuals, they might be necessary for the adaptation and survival of the species, especially in slow-reproducing organisms. Otherwise, it is hard to imagine how the accumulation of tens of thousands of transposons near essential genes could be tolerated over millions of years. Our analysis show that these TEs may influence development from an early formative stage. Thus expansion of different TEs that contribute to the rewiring of developmental pathways may facilitate speciation and adaptation.

#### Gene regulation by transposons

TEs can be involved in epigenetic regulation, as they can recruit the silencing machinery^19,20,36^. Many examples have been reported that retrotransposons can serve as TF binding sites to promote nearby gene expression. In addition to LTRs, which contain promoters that can be used to drive expression of nearby genes^32^, LINEs elements contain an antisense promoter that can be used by nearby genes^33,111,112^. MER20, a DNA transposon, was found to contribute to pregnancy-related gene network and its evolution^46^. In ESCs, Kunarso et al ^34^ reported that about 25% of POU5F1, NANOG and CTCF binding sites are provided by TEs. Thus TEs is important in the regulatory network of ESCs^37^. More importantly, retroviral activity was found to be a hallmark of pluripotency^26,27,31,113^. ERV-derived LTR elements may be contributing to the gene regulatory network of innate immunity^114^. Adding to these results, this study systematically investigated the correlation of TEs and the genomics reprogramming in PD. We show that TEs maybe play a more profound role than previously thought, affecting thousands of genes. We also provide some evidence for the potential role of SINEs in activating housekeeping and ESC-specific genes.

### Possible mechanisms of B1 and Alu in gene regulation

Mouse B1 and human Alu elements originated from 7SL RNA^64^ and contain RNA polymerase III promoters. The A-box and B-box included in Alu sequences are bound by a multi-subunit transcription factor TFIIIC, to form the Pol III complex. Although some microRNAs are shown to be transcribed by Pol III using upstream Alus^115^, it is unlikely that Pol III would produce thousands of essential genes. This would predict a strand-specific correlation and alternative TSS’, similar to LTRs. Alu family repeats also contain binding sites for many factors associated with RNA Pol II ^116,117^, including p53^118^, retinoic acid receptors ^119,120^, YY1^121^, PIT2^122^, *etc*. Our analysis shows that B1 elements also contain TFBS’ for Obox family proteins, especially Obox3. These TFs may act upon Alu elements to drive gene expression.

In addition, it is well documented that TFIIIC, without the rest of the Pol III apparatus, has the so-called extra-transcriptional effects (ETC) ranging from nucleosome positioning, genome organization, and direct effect on Pol II transcription^123,124^. Alu provides the majority of the TFIIIC binding sites in humans and mice^125^. B2 elements are also enriched with CTCF binding motifs defined by Chip-Seq^9,19^.

Some human Alus serve as estrogen receptor (ER)-dependent enhancers for BRCA1^126^. Su *et al.* found that human Alu elements in the proximal upstream region are more conserved and show many properties of enhancers^127^. Indeed, similar to enhancer RNAs (eRNAs)^128^, transcription of Alu transposons may boost the expression of downstream genes through an enhancer mechanism specific to embryonic cells. Further study is needed to investigate these possible mechanisms.

Transposons near genes should be treated more like regulatory elements. In order for a single TF to regulate a large number of genes, it can evolve to take advantage of existing mobile elements. This is more likely than the scenario where hundreds of genomic loci converge to the binding motif of an existing TF, which is especially true for highly specific motifs such as that of CTCF^129^.

Guided by biological curiosity, exploratory analysis of the single-cell RNA-seq and related data yields many actionable insights. One of the surprising observations is that genes with similar expression patterns in early embryogenesis share specific transposons in their promoters. During ZGA, while LTRs are linked to transient, forceful and early induction of several hundred genes, SINE elements are associated with the upregulation of thousands of essential genes. The machinery that transcribes retro-transposons may also be used to establish the expression landscape of early embryos. This study also demonstrates the power of single-cell RNA-seq, especially when applied to the study of normal developmental processes.

## Methods

Raw data for the single-cell RNA-seq were downloaded from NCBI’s Short Read Archive with accession number PRJNA195938 using the fastq-dump program of SRAtools suite. FastQC was used for the initial quality check^130^. Trimming of sequences was carried out using cutadapt^131^. Mouse genome sequence (GCRm38) and annotation were downloaded from ENSEMBL using the biomaRt^82^ package on Bioconductor^132^. We used the Tophat and cufflinks programs^48^ to map and quantify gene expression. Translation starting sites (TSSs) of genes were defined as the TSS of the highest expressed transcript isoforms across all the samples in this study. Read mappings of RNA-seq data were generated by Integrative Genomics Viewer (IGV)^133^. ALEA software^134^ was used to map the reads to maternal and paternal alleles based on single nucleotide polymorphisms (SNPs) derived from genome sequences of the two strains^135^. In order to estimate retrotransposon expression, we re-mapped reads using STAR^136^ to allow more multiple-mapped reads using the following parameters: STAR --outFilterMultimapNmax 100 --winAnchorMultimapNmax 100 --outSAMmultNmax 100 --outSAMtype BAM SortedByCoordinate --outFilterMismatchNmax 3. The expression levels of retrotransposons were calculated using TEtranscripts^50^, which was specially designed to estimate both gene and TE abundances by using an additional index of TEs based on UCSC repeatMasker files. The parameters used are: TEtranscripts --format BAM --mode multi --GTF genes.gtf --TE mm10_rmsk_TE.gtf -i 2 --stranded no. A supplementary file gives all commands used in sequence analyses.

The key features in our TF binding analysis are: 1) use of RNA-seq data for TSS location, 2) use the highest score as an indicator to avoid arbitrary cutoff in deciding TF binding, 3) rank genes by fold-change to avoid cutoff in gene clusters, and 4) filter and prioritize using the expression pattern of TF genes.

In the clustering analysis, genes with expression levels less than 5 FPKM across all samples were eliminated from analysis. The remaining genes were sorted by standard deviation and the top 12,000 were selected. Cluster 3.0^137^ and Java TreeView^138^ were used for hierarchical clustering and visualization.

To analyze the expression profiles of transposons, the reads were mapped to consensus sequences of repeats in RepBase (version 19.04)^54^ using Bowtie^139^. After correction for library size, total mapped reads per transposon were used as an approximate indicator of relative expression levels.

## Acknowledgements

This research used computers managed by Administrative and Research Computing at South Dakota State University. The work is partially supported by National Institute of Health (GM083226), the National Science Foundation/EPSCoR Grant Number IIA-1355423, and the State of South Dakota.

## Conflict of interest

None.

